# Integrating Literature Evidence to Enhance CRISPR-based Target Screening for Hand-Foot Syndrome

**DOI:** 10.1101/2025.03.09.642163

**Authors:** Qiqi Yang, Bingxue Yang, Junkang An, Dazhao Lv, Shuyue Xu, Qing You, Jie Luo, Shiyi Zhang

## Abstract

The experimental high-throughput screening (HTS) methods, exemplified by CRISPR-based screening techniques, have revolutionized target identification in drug discovery. However, such screens frequently yield extensive, often unrelated target lists necessitating costly and time-intensive experimental evaluation and validation. To address this challenge, we propose a dual-filter strategy that integrates literature-mined targets with CRISPR/Cas9 screening outputs, systematically prioritizing the most credible candidates and thereby reducing the experimental validation burden and increasing success rate. To validate this strategy, we applied it with hand-foot syndrome (HFS), a clinically challenging side effect induced by fluoropyrimidine treatment. We identified ATF4 as a key regulator of 5-fluorouracil (5-FU) toxicity in the skin and revealed forskolin as a potential therapeutic agent of HFS through the strategy. Mechanistically, forskolin triggers MEK/ERK-dependent ATF4 induction, subsequently driving 5-FU detoxification via the ATF4-mediated eIF2α/IκB signaling pathway. Our findings demonstrate that this dual-filter strategy could notably accelerate drug discovery by reducing experimental validation burden after target screening.

## Introduction

Identifying targets for diseases is a critical step in the drug development process. Among the established approaches, experimental high-throughput screening (HTS) methods aim to rapidly extract potential targets from large-scale datasets, including functional genomics, genome-wide association studies, and omics analysis. Genome-wide CRISPR/Cas9 screening technology is widely recognized for its powerful ability to identify causal relationships between genetic phenotypes(*1*). Currently, this target screening method has been widely used in various research fields, including cancer(*2*), antibiotic discovery(*3,4*), viral infections(*5*) and diabetes(*6*), and greatly improved drug development efficiency. However, despite the advantages, CRISPR/Cas9 screening often exhibits high false-positive rates and lacks reproducibility(*7*). This makes extensive experimental verification necessary for the identification of the most promising candidate targets (*8*), the process that is both costly and time-consuming, consequently augmenting the overall burden of drug development. Therefore, developing approaches to refine screening results and to reduce the workload associated with target validation is critical for improving the efficiency of the drug discovery process.

To improve the success rate of target verification, several approaches have been explored to enhance the accuracy of target identification. Narrowing the scope of target screening through functional classification of genes is a common method to filter CRISPR/Cas9 screening outputs. Some studies classify genes by excluding irrelevant genes like housekeeping genes(*9*) from screening results or employing specialized libraries before experimental screening(*1,10*). However, since the classifying process is typically based on existing experience-dependent functional annotations, these approaches are applicable only in specific research contexts where functional classification is available. Other studies first identify relevant genes with specific function through HTS like omics studies and subsequently use these genes to classify screening outputs(*1,11*). Compared to the former approach, this method offers greater flexibility in application. However, as it still originates from large-scale experimental data, this approach is often subject to the same limitations as CRISPR/Cas9 sreening.

Notably, SkywalkR app was built to simulate the manual ranking process of experts by integrating diverse types of evidence, including preclinical, clinical, literature, and gene essentiality data(*12*). However, this multi-objective ranking approach, with the absence of mature technology for integrating heterogeneous datasets, may lead to the over-ranking of genes with high scores in a single criterion and also result in challenging to identify the most optimal target due to the insufficient differentiation between high-score genes. Inspired by the designation and limitation of SkywalkR, we provided a strategy with more applicable flexibility and less limitation to enhance efficacy of target identification.

Scientific literature constitutes a key repository of experimentally validated therapeutic targets across diverse disease domains. Integrating literature data with CRISPR/Cas9 screening results offers an alternative filtering strategy to reduce the workload of researchers by enabling the substitution of labor-intensive experimental validation with computational validation. Therefore, our target screening strategy aims to employ literature-based screening as the second filter to prioritize CRISPR/Cas9 screening outputs.

To evaluate the potential of our strategy in enhancing the efficiency of target identification, we used hand-foot syndrome (HFS), a common side effect of fluoropyrimidine chemotherapy, as a study case. Capecitabine, a prodrug of 5- fluorouracil (5-FU), is currently a major chemotherapy agent causing HFS (*13*). The administration of capecitabine induces a situation where approximately 40% of patients necessitate dose modification (*14*), with the primary causative factor being HFS (*15*). Although as a dose-limiting side effect, HFS lacks other clinical standard and effective management, highlighting the urgent need to discover additional candidate drugs.

The pathogenesis of HFS is believed to result from the accumulation of 5-FU in the hands and feet, leading to keratinocyte damage (*16*). Thymidine diacetate, a prodrug of thymidine, can counteract the inhibitory effect of 5-FU on TS by supplementing thymidine. Local administration of this drug has demonstrated efficacy in relieving HFS(*17*). Uridine can mitigate 5-FU toxicity by increasing the intracellular UTP pool to compete with FUTP, the latter is one of the activated forms of 5-FU, which damages RNA by incorporating into RNA (*18*). 10% uridine ointment has been shown to relieve HFS symptoms (*19,20*). An inhibitor of TP also exhibit efficacy in relieve HFS by inhibiting the transform of 5-DFUR into 5-FU (*21*). ALL above strategies detoxify 5- FU by targeting the 5-FU mechanism but ignored the role of cell response to 5-FU. Therefore, this study aims to discover therapeutic candidates for HFS through systematic investigation of molecular targets modulating 5-FU-induced cutaneous toxicity.

Genome-wide CRISPR/Cas9 screening technology is commonly employed to identify targets for combination therapies that enhance the sensitivity or reduce the resistance of chemotherapy(*22,23*). Through integrating literature-based screening with genome-wide CRISPR/Cas9 screening, we identified ATF4 as a promising target for modulating 5-FU toxicity in the skin. Subsequent efficacy validation of forskolin, an ATF4 inducer, demonstrated its ability to detoxify 5-FU and alleviate HFS. Further exploration of the biological mechanisms underlying forskolin’s action also confirmed the function of ATF4 in regulating 5-FU toxicity.

In conclusion, our study presents an effective target screening strategy for drug discovery using HFS as an application case.

## Results

### 1. Genome-wide CRISPR/Cas9 screening identified targets regulating 5-FU toxicity in HaCaT

We conducted genome-wide CRISPR/Cas9 screening on HaCaT cells under 5-FU treatment to identify modulators of 5-FU toxicity in keratinocytes. We selected the Human GeCKOv2 Library A, which targets 19,050 genes, with three sgRNAs per gene, and packaged them into lentivirus. The exogenous DNA carrying sgRNAs sequence in lentivirus was then transduced to HaCaT cells expressing Cas9 protein at a multiplicity of infection (MOI) of 0.3. Successfully transduced cells with sgRNAs were selected and amplified with puromycin. The pool of library-transduced cells was divided into three groups: a control group treated with PBS, and two treatment groups exposed to 5- FU at IC50 and IC99 concentrations respectively. Each group was cultured under its respective conditions for 10 days. At the end of experimental screening, the genomic DNA in all groups were sequenced using next-generation sequencing (NGS). Each sgRNA count hit by NGS represents a single cell with a gene knocked out. Data were analyzed by MAGeCK (*24*). The normalized sgRNA counts and robust rank aggregation (RRA)scores are available in separate files (**tables S1 and S2**). Finally, we obtained two target libraries, Library-IC50 and Library-IC99 **(Fig. 1A).**

**Fig. 1.**
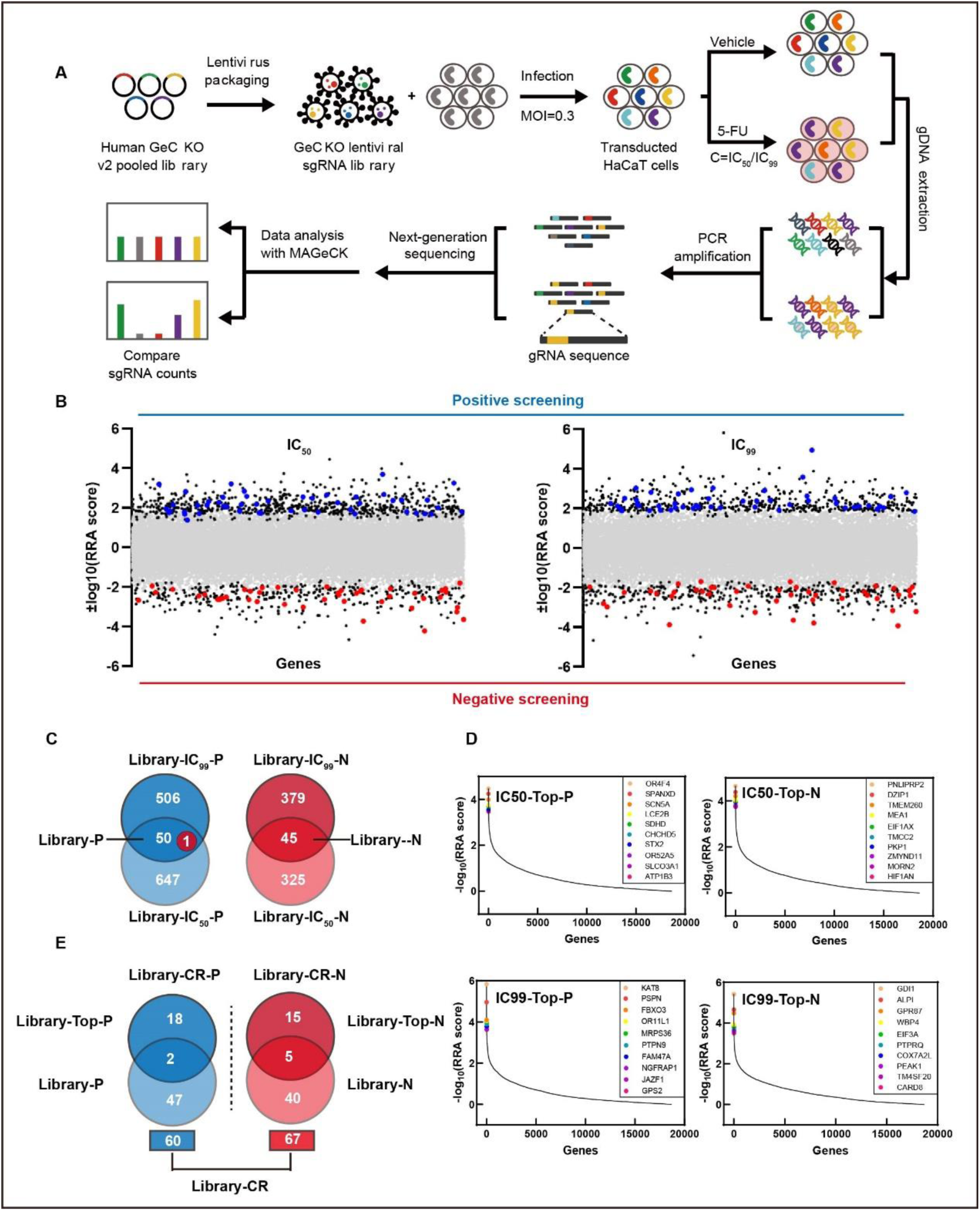
**Genome-wide CRISPR/Cas9 screening results**. (A) Experimental workflow of genome-wide CRISPR/Cas9 screening. (B) RRA score distribution of Library-N and Library-P. (C) Definitions of Library-N and Library-P. (D) Top 10 targets screened by genome-wide CRISPR/Cas9. (E) Definition of Library-CR.

For either of the two libraries, we distinguished negative and positive screening results by ranking the RRA score. In Library-IC50, targets with p-value < 0.02 were distinguished into Library-IC50-N (negative screening) and Library-IC50-P (positive screening); in Library-IC99, we also obtained Library-IC99-N and Library-IC99-P. Targets present in both Library-IC50-P and Library-IC99-P, but absent from all other libraries, were classified into the Library-P group, consisting of 49 genes marked with green dots **(Fig. 1, B and C).** In the same way, Library-N contains 45 genes marked with red dots **(Fig. 1, B and C).** In addition, we extracted the top 10 targets from the negative or positive screening of Library-IC50 and Library-IC99 to form Library-Top-N and Library-Top-P, respectively **(Fig. 1, D and E)**. Ultimately, we consolidated two libraries from CRISPR/Cas9 screening: Library-CR-N, comprising 60 targets, and Library-CR-P, comprising 67 targets **(Fig. 1E)**. Combining the two libraries, we established Library-CR, which includes 127 targets potentially influencing 5-FU toxicity in keratinocytes. All target libraries identified in this study are available in a separate file **(table S3)**.

### 2. Text mining screening identified targets regulating 5-FU toxicity in the skin

Our genome-wide CRISPR/Cas9 screening revealed 127 putative genetic modulators associated with 5-FU toxicity, all requiring functional validation through experimental characterization. We conducted a literature screening via text mining technology to identify literature-supported targets related to 5-FU toxicity regulation in order to alleviate the burden of experimental verification by secondly filtering the CRISPR/Cas9 screening outputs.

However, the limited availability of literature in the field of fluoropyrimidine-induced HFS constrained comprehensive text mining. To overcome this challenge, we devised an approach to augment available literature by incorporating other research domains related to 5-FU beyond the scope of HFS. The applied 5-FU dosage correlates with its anti-tumor efficacy and the severity of HFS in a similar way. Thus, the genes regulating 5-FU toxicity in cancer cells may have the same impact on the effects of 5-FU in keratinocytes, prompting us to expand the available literature from the domain of HFS to anti-tumor research. In order to leverage available literature from anti-tumor research, we extracted two key parameters pertinent to HFS: the chemotherapeutic drug “5-FU” and the affected tissue “skin”, and screened targets from the two aspects separately.

The literature-based targets screening process is shown in **Fig. 2A**. We first conducted a search using the keywords “5-FU AND (resistance OR sensitivity) AND (expression OR silence OR knockdown)” in the PubMed database for the information in title and abstract. We used the “Tokenizer” program to segment the text from them (*25*), generating a list of individual words with information stored in a vector. The reference gene list including gene symbols and gene aliases is provided by the National Center for Biotechnology Information (NCBI) **(table S4)** with information stored in another vector. During the “Extractor” process, we calculated vector similarity between the two vector separately from “Tokenizer” and from “NCBI”. This dual-vector architecture enabled precise semantic matching between biomedical literature content and established gene nomenclature. Ultimately, we extracted the genes reported to be associated with 5-FU toxicity.

**Fig. 2.**
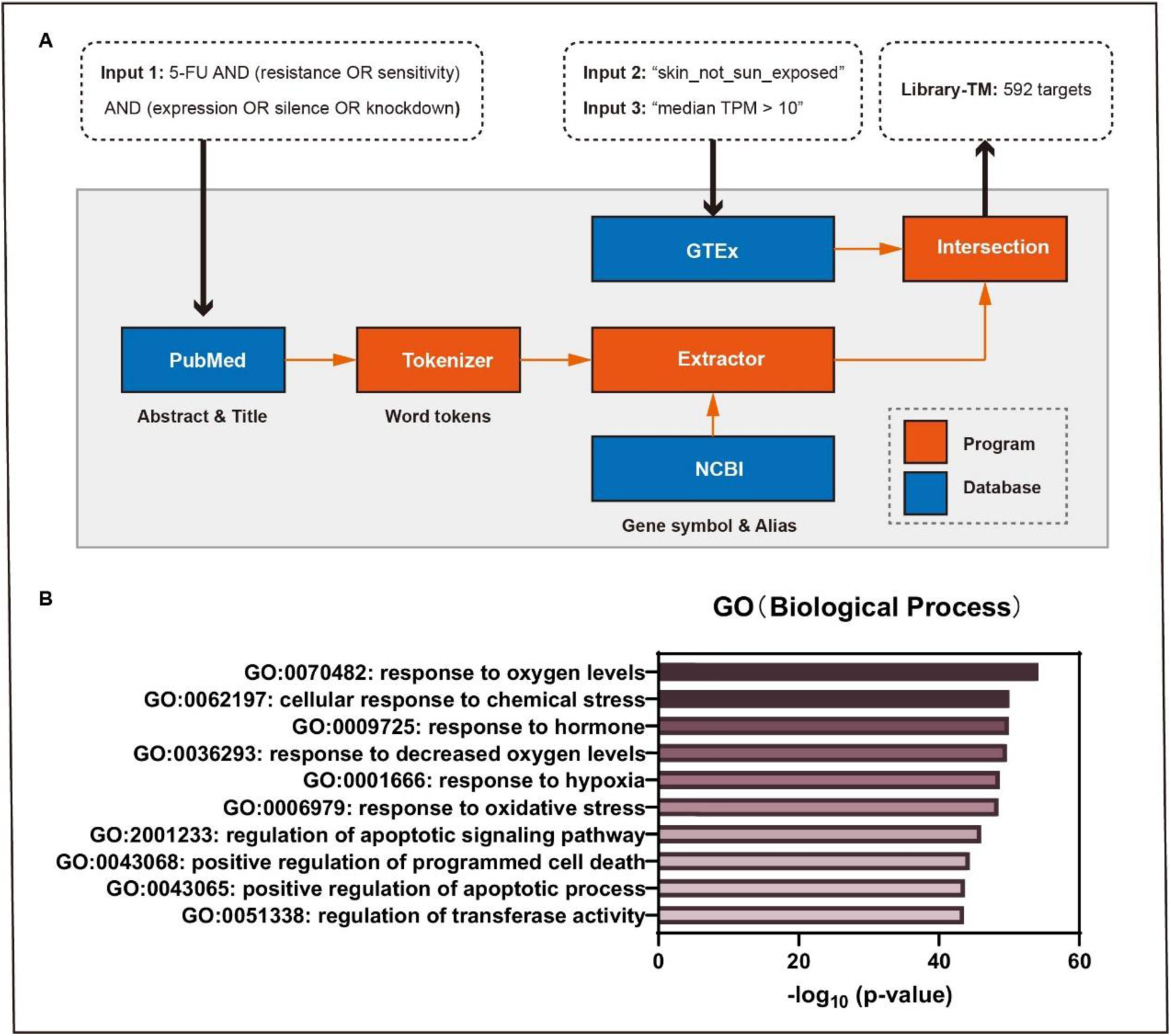
Text mining screening results. (A) Workflow of text mining screening. (B) GO enrichment analysis of Library-TM.

Since the genes identified by the “Extractor” are reported in different types of cells and tissues, to ensure the screened targets play a crucial role in skin-limited side effects, we extracted distinct targets highly expressed in the skin by leveraging the GTEx database, with the median TPM > 10. Finally, the intersection of targets from literature screening and GTEx database yielded the Library-TM, a target library based on text mining, consisting of 592 genes **(Fig. 2A and table S5).**

GO enrichment analysis of Library-TM revealed that most enriched gene function terms (top 6 terms) are associated with “response to stimulus” **(Fig. 2B)**. This suggests that the biological processes involved in the cellular response to stimulus may regulate 5- FU toxicity.

### 3. ATF4 was identified through integrating literature evidence with CRISPR/Cas9 screening

We applied the 592 targets identified through literature-based screening to refine the 127 targets from the CRISPR/Cas9 screening by Library-CR and Library-TM. We identified two genes through the dual-filter strategy, *ATF4* and *MAPK14* **(Fig. 3A)**.

**Fig. 3.**
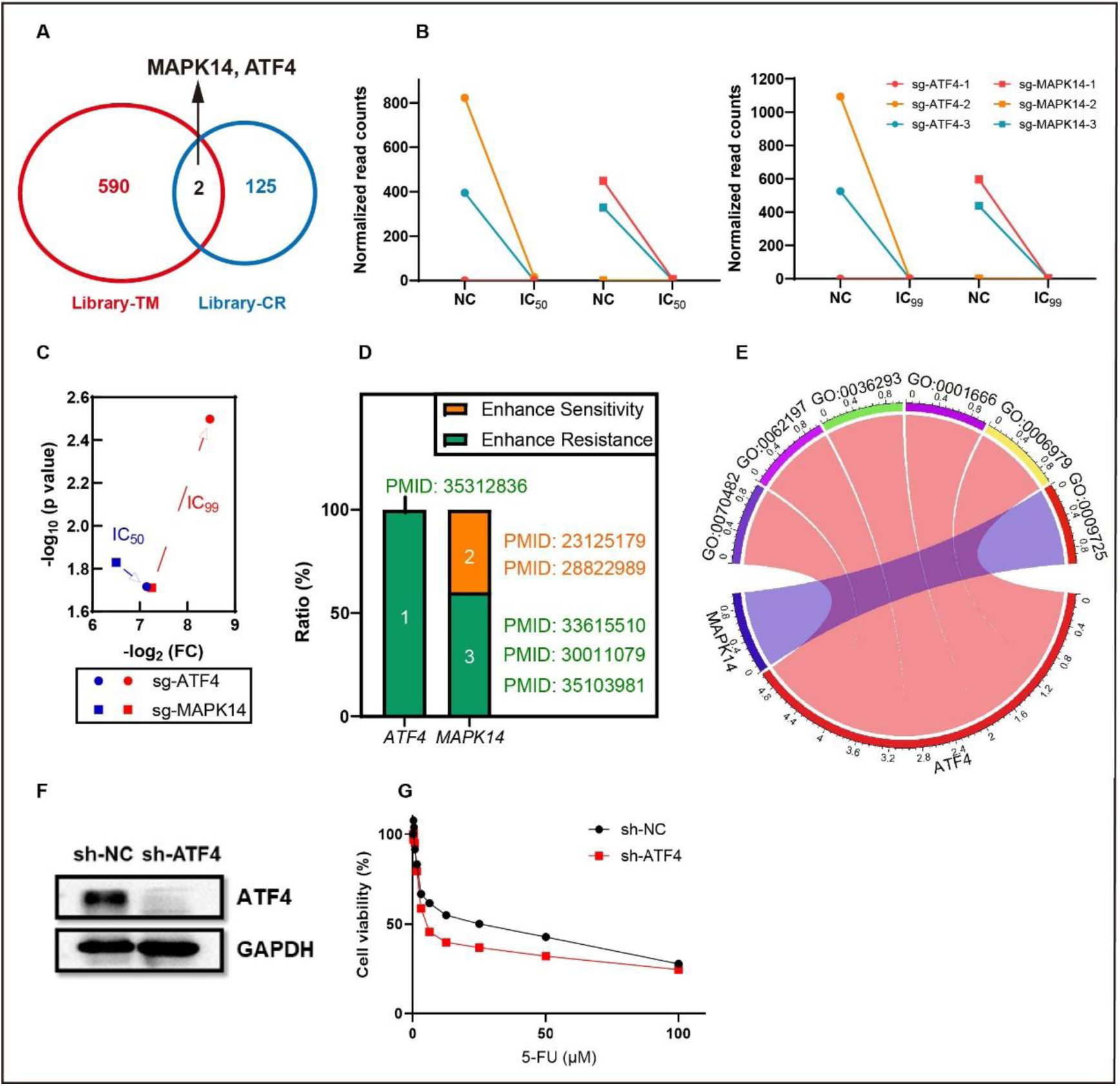
Targets identification process. (A) *ATF4* and *MAPK14* are identified through combining CRISPR/Cas9 and text mining screening. (B) *ATF4* knockout has a stronger influence on normalized sgRNA counts. (C) Comparison of the changes in sgRNA counts and its statistical significance (D) The reported regulatory ability on 5-FU toxicity of *MAPK14* shows poor consistency. (E) The occurrence frequencies of *ATF4* and *MAPK14* in stress-related GO terms. (F) ATF4 protein expression in the *ATF4* knockdown cell line. (G) The impact of *ATF4* knockdown on 5-FU IC_50_

In the CRISPR/Cas9 screening, *ATF4* knockout resulted in a more pronounced change in normalized sgRNA counts than *MAPK14* knockout **(Fig. 3B)**. *ATF4* knockout also exhibited greater potential in regulating 5-FU toxicity with supporting evidence both in statistical significance and fold change **(Fig. 3C)**. Additionally, we evaluated the performance of *ATF4* and *MAPK14* in literature screening. *ATF4* was reported to enhance 5-FU resistance, whereas *MAPK14* was noted to exert a contradictory capability in regulating 5-FU toxicity. Two in five articles reported its function in enhancing 5-FU sensitivity, which is inconsistent with the CRISPR/Cas9 screening results **(Fig. 3D)**. The regulation of 5-FU toxicity was related to “response to stimulus” as indicated by the top six GO enrichment terms of Library-TM. *ATF4* appeared in five terms, while *MAPK14* in only one term, suggesting a stronger functional link between ATF4 and 5-FU toxicity **(Fig. 3E)**. Therefore, as indicated by the CRISPR/Cas9 screening results, *ATF4* holds greater potential than *MAPK14* for modulating 5-FU toxicity.

Our findings confirmed that knockdown and knockout of *ATF4* enhances 5-FU sensitivity in HaCaT cells **(Fig. 3, F and G, and fig. S1)**, supporting the role of ATF4 in regulating 5-FU toxicity as suggested by both CRISPR/Cas9 and text mining analyses. Finally, *ATF4* is identified as a candidate gene for modulating 5-FU toxicity.

### 4. Forskolin reverses 5-FU toxicity in HaCaT as an ATF4 inducer

*ATF4* knockdown was indicated to enhance 5-FU toxicity, driving us to discover the agent detoxifying 5-FU through increasing ATF4 expression. Forskolin has been reported to induce ATF4 expression (*26*), which was also verified in our experiments **(Fig. 4A and fig. S2A).** Therefore, forskolin was expected to reverse 5-FU toxicity by upregulating ATF4.

**Fig. 4.**
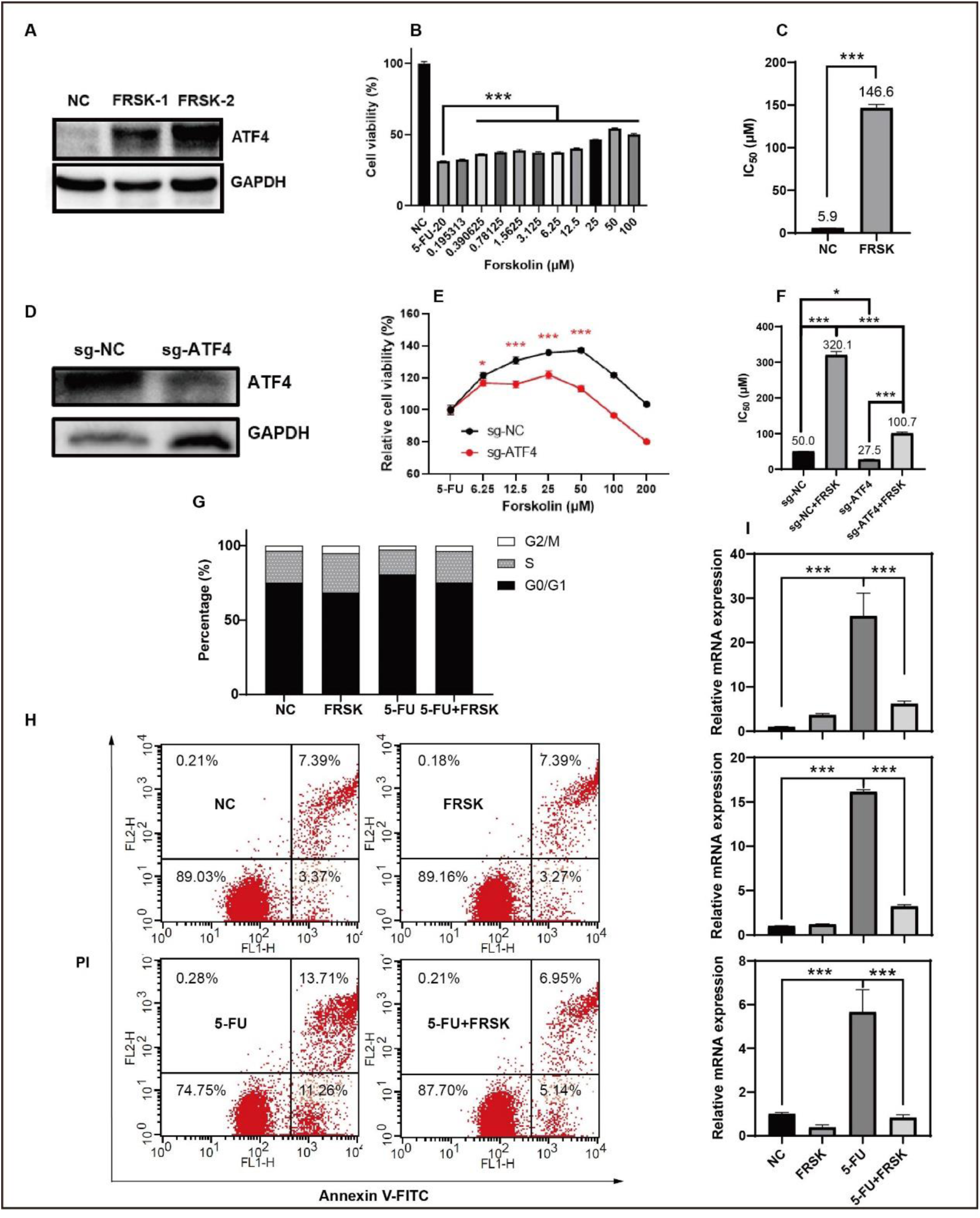
The efficacy of forskolin in detoxifying 5-FU evaluated on HaCaT. (A) Forskolin significantly induced ATF4 expression. (B) Forskolin alleviated 5-FU-induced cytotoxicity, and (C) increased 5-FU IC_50_. (D), (E), (F) *ATF4* knockout impaired forskolin’s ability to reverse 5-FU toxicity (G) cell cycle arrest, (H) cell apoptosis, and (I) the release of typical pro-inflammatory cytokines.

The CCK-8 assay and LDH release assay demonstrated that forskolin treatment reversed 5-FU-induced toxicity in HaCaT cells in a dose-dependent manner **(Fig. 4B and fig. S2B)** and significantly increased the IC50 of 5-FU in HaCaT **(Fig. 4C)**. As a plant extract, forskolin may exert its effects through various unknown pathways. This drives us to explore the role of ATF4 in forskolin-induced 5-FU detoxification. Our results showed that *ATF4* knockout diminished the dose-dependent protection of forskolin **(Fig. 4, D and E)** and impaired the increase of 5-FU IC50 induced by forskolin **(Fig. 4, D and F).** This indicates that ATF4 contributes partially to the detoxification effect of forskolin. The remaining forskolin’s effects may derive from other signal pathways downstream of the MEK/ERK activation, but not the induction of ATF4 **(fig. S2C)**.

5-FU is known to induce cell or tissue damage by blocking the cell cycle, promoting apoptosis, and increasing the release of pro-inflammatory factors(*27–29*). According to our results, forskolin was found to counteract 5-FU’s effects on cell cycle arrest **(Fig. 4G)**, inducing apoptosis **(Fig. 4H)**, and upregulating the expression of classical pro-inflammatory factors (IL-6, IL-1β, TNF-α) **(Fig. 4I)**, colony-stimulating factor (GSF- 2), and chemokines (CXCL-1) **(fig. S2, D and E).** These findings suggest that forskolin has the potential to act as a detoxifying agent against 5-FU.

### 5. Forskolin reverses HFS induced by capecitabine in vivo

In vitro findings demonstrated the ability of forskolin to rescue 5-FU toxicity in HaCaT cells, reflecting its potential to alleviate HFS. We formulated a gel containing 10% forskolin. After daily administration of capecitabine, forskolin-containing gel (CAP- FRSK) or vehicle gel (CAP-Vehicle) was applied topically to the hind paws of rats. The progression of HFS was monitored and assessed using a disease grading system: Grade 1 indicated mild dryness, while Grade 2 represented severe dryness, desquamation, or dryness with redness and swelling. By day 35, as approximately 90% of rats in the CAP-Vehicle group (HFS model group) exhibited Grade 2 HFS, we ended the experiment.

Notably, the severity of HFS in the CAP-FRSK group was significantly lower than CAP-Vehicle group **(Fig. 5A).** The incidence of severe HFS like Grade 2 HFS during chemotherapy is crucial for the clinical decision of dose modification (*30*). Therefore, we analyzed the effect of forskolin on the incidence of Grade 2 HFS. Our analysis revealed a lower daily proportion of Grade 2 HFS in the CAP-FRSK group, indicating a slower progression of symptoms **(Fig. 5B)**. At the end of the experiment, the incidence rate for Grade 2 HFS during the whole experiment was 91% (n=12) for CAP-Vehicle and only 25% (n=12, one death on day 26) for CAP-FRSK. The images of rats’ hind paws at the termination showed that HFS in the CAP-Vehicle group presented severe desquamation and redness, while forskolin treatment resulted in milder symptoms **(Fig. 5C).** Moreover, pain sensitivity was reduced in the CAP-FRSK group compared to the CAP-Vehicle group **(Fig. 5D).**

**Fig. 5.**
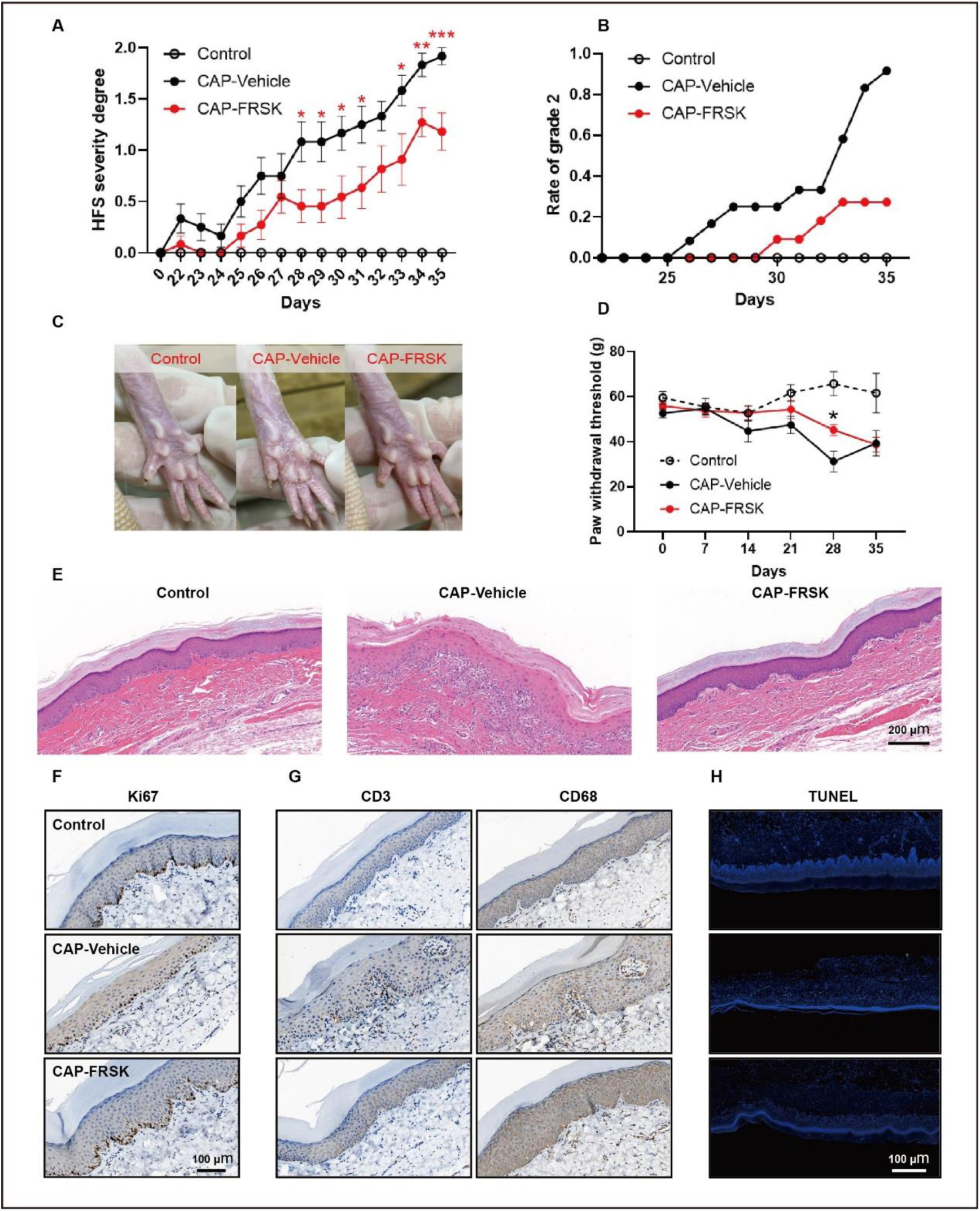
The efficacy of forskolin in alleviating HFS evaluated on rats. (A) Forskolin relieves the severity of HFS in rats. (B) Forskolin reduced the rate of grade 2 HFS on rats. (C) Representative images of HFS symptoms on the rats’ hind paws. (D) Forskolin elevated paw withdrawal threshold. (E) HE stains images of the full-thickness skin of the rats’ hind paws. (F), (G) IHC stains of Ki6s7, CD3 and CD68. (H) The TUNEL stains images of the rats’ hind paw skin.

HE stains of the hind paw skin revealed that the HFS model group (CAP-Vehicle) exhibited dyskeratosis (premature keratinization that did not reach the stratum corneum), disordered basal layer structure, incomplete keratinization (presence of stratum corneum cell nuclei and lack of granular layer), and immune cell infiltration. These pathological changes were alleviated in the CAP-FRSK group **(Fig. 5E)**. IHC staining of Ki67 indicated that forskolin alleviated the inhibition of keratinocyte proliferation induced by capecitabine **(Fig. 5F)**. In addition, forskolin might also exert anti-inflammatory effects by inhibiting lymphocyte and macrophage infiltration in the epidermis **(Fig. 5G)**. Furthermore, TUNEL assay indicated that forskolin could alleviate capecitabine-induced keratinocyte apoptosis **(Fig. 5H).**

In the Balb/c mice model, forskolin also attenuated HFS progression, with a Grade 2 incidence of 50% (n=10, one death on day 11) in the CAP-Vehicle group compared to 0% (n=10) in the CAP-FRSK group at the end of the experiment. However, spontaneous HFS remission was observed over time **(fig. S3, A and B)**. Forskolin treatment substantially alleviated HFS symptoms on Balb/c mice, including scaling, redness, swelling, and pain **(fig. S3, C and D).**

In conclusion, these findings indicate that forskolin has considerable potential in mitigating capecitabine-induced HFS.

### 6. Forskolin mitigates 5-FU effects via the ATF4-eIF2α-IκB signaling pathway

Our in vivo and in vitro experiments demonstrated that forskolin reverses 5-FU toxicity and alleviates HFS induced by capecitabine. However, the precise mechanism by which forskolin facilitates5-FU detoxification through ATF4 induction remains unclear.

ATF4 has two classical functions (*31*). First, as a key downstream effector of the integrated stress response (ISR), ATF4 enhances the transcription of various stress-response proteins to respond to the upregulated p-eIF2α. Second, as a negative feedback regulator of ISR, ATF4 dephosphorylates p-eIF2α by upregulating GADD34.

Studies have shown (*26*) that forskolin-induced ATF4 could dephosphorylate p-eIF2α, which is also verified on HaCaT in our results **(Fig. 6, A and B)**. In addition, forskolin treatment failed to increase the transcription of most genes downstream of ATF4 except *GRP78* and *P16*, but the protein levels of both genes remained unchanged **(fig. S4)**. This evidence demonstrated that forskolin-induced ATF4 predominantly mediates its therapeutic effects through the second pathway rather than the first one. Several studies have shown that inhibition of p-eIF2α exhibits a cellular protective effect (*32–34*). Therefore, the aforementioned evidence suggests that forskolin may reverse 5-FU toxicity through dephosphorylating p-eIF2α.

**Fig. 6.**
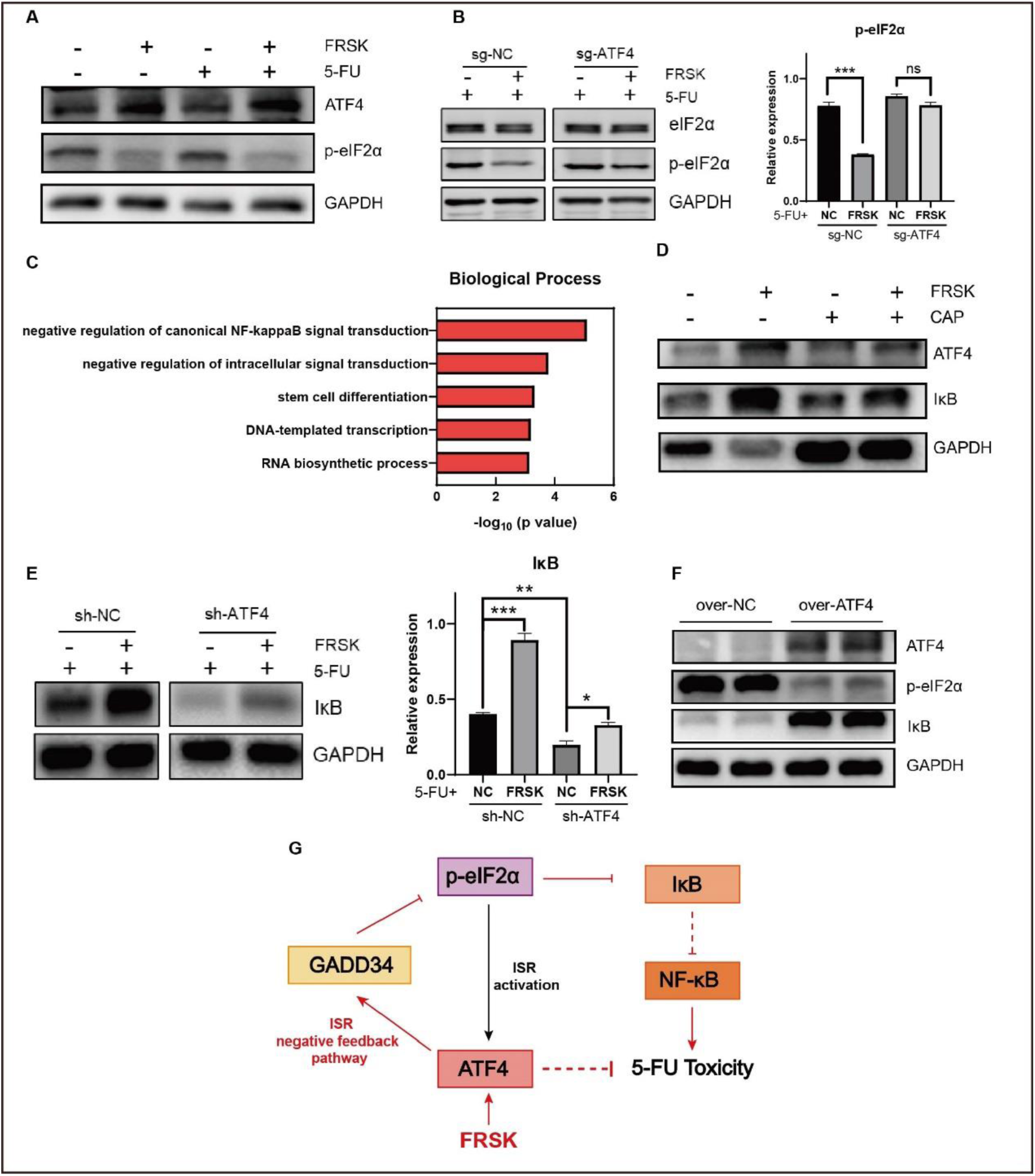
The biological mechanism underlying forskolin action. (A) 12 h forskolin treatment downregulated p-eIF2α in HaCaT. (B) *ATF4* knockout impaired forskolin’s ability to downregulate p- eIF2α in HaCaT. (C) Genes in Library-CR-N were enriched in negative regulation of NF-κB pathway. (D) forskolin treatment upregulated IκB expression in rats’ hind paw skin. (E) *ATF4* knockdown impaired forskolin’s ability to upregulate IκB in HaCaT.(F) *ATF4* overexpression replicated the effect of forskolin on p-eIF2α and IκB expression in HaCaT. (G) The possible biological mechanism of forskolin to detoxify 5-FU.

To investigate the role of ATF4/eIF2α in mediating 5-FU detoxification, we performed GO enrichment analysis on Library-CR-N where ATF4 was identified. This analysis revealed enrichment in the “negative regulation of NF-κB pathway” **(Fig. 6C)**, suggesting that inhibition of the NF-κB pathway may reduce 5-FU toxicity to HaCaT cells. In addition, RNA-seq of *ATF4*-knockdown HaCaT cells confirmed that differentially expressed genes were enriched in the NF-κB pathway **(fig. S5)**, supporting the hypothesis that ATF4 regulates the NF-κB pathway in HaCaT cells to reverse 5-FU toxicity. Furthermore, elevated p-eIF2α levels are known to induce inflammation by activating NF-κB (*35*), which is connected with impaired IκB expression (*36*). This evidence suggests us to investigate the effects of forskolin on the ATF4-eIF2α-IκB pathway. Forskolin was found to increase IκB expression in rat skin tissues **(Fig. 6D)**. At the same time, the knockdown of *ATF4* attenuated the upregulation of IκB induced by forskolin **(Fig. 6E)**, indicating that forskolin’s effect on IκB could be mediated through ATF4. Additionally, overexpression of *ATF4* successfully replicated forskolin’s effects, including downregulated p-eIF2α and upregulated IκB **(Fig. 6F)**.

In summary, forskolin-induced ATF4 reduces p-eIF2α level via the negative feedback pathway of ISR, leading to increased IκB expression **(Fig. 6G)**. Forskolin’s effects on IκB were believed to derive from enhanced global translation (*37*). Upregulated IκB may subsequently regulate survival, proliferation, inflammation, and other phenotypes by inhibiting the NF-κB pathway.

### 7. Topical forskolin treatment does not affect the anti-tumor efficacy of capecitabine

Although forskolin is designed to be administrated topically, as an adjuvant drug for chemotherapy, it necessitates concomitant administration with capecitabine. This may compromise the anti-tumor efficacy of capecitabine by triggering adverse drug-drug interactions. Therefore, we investigated the impact of local forskolin treatment on the anti-tumor efficacy of capecitabine.

We established a subcutaneous tumor-bearing nude mouse model using the human colorectal cancer cell line HCT116. When tumor volumes reached approximately 150 mm³, capecitabine treatment was initiated concurrently with the topical application of either forskolin ointment (CAP-FRSK) or the corresponding vehicle (CAP-Vehicle). Throughout the study, no significant differences in body weight were observed between the two groups **(Fig. 7A)**, suggesting that topical forskolin might not cause systemic metabolic changes.

**Fig. 7.**
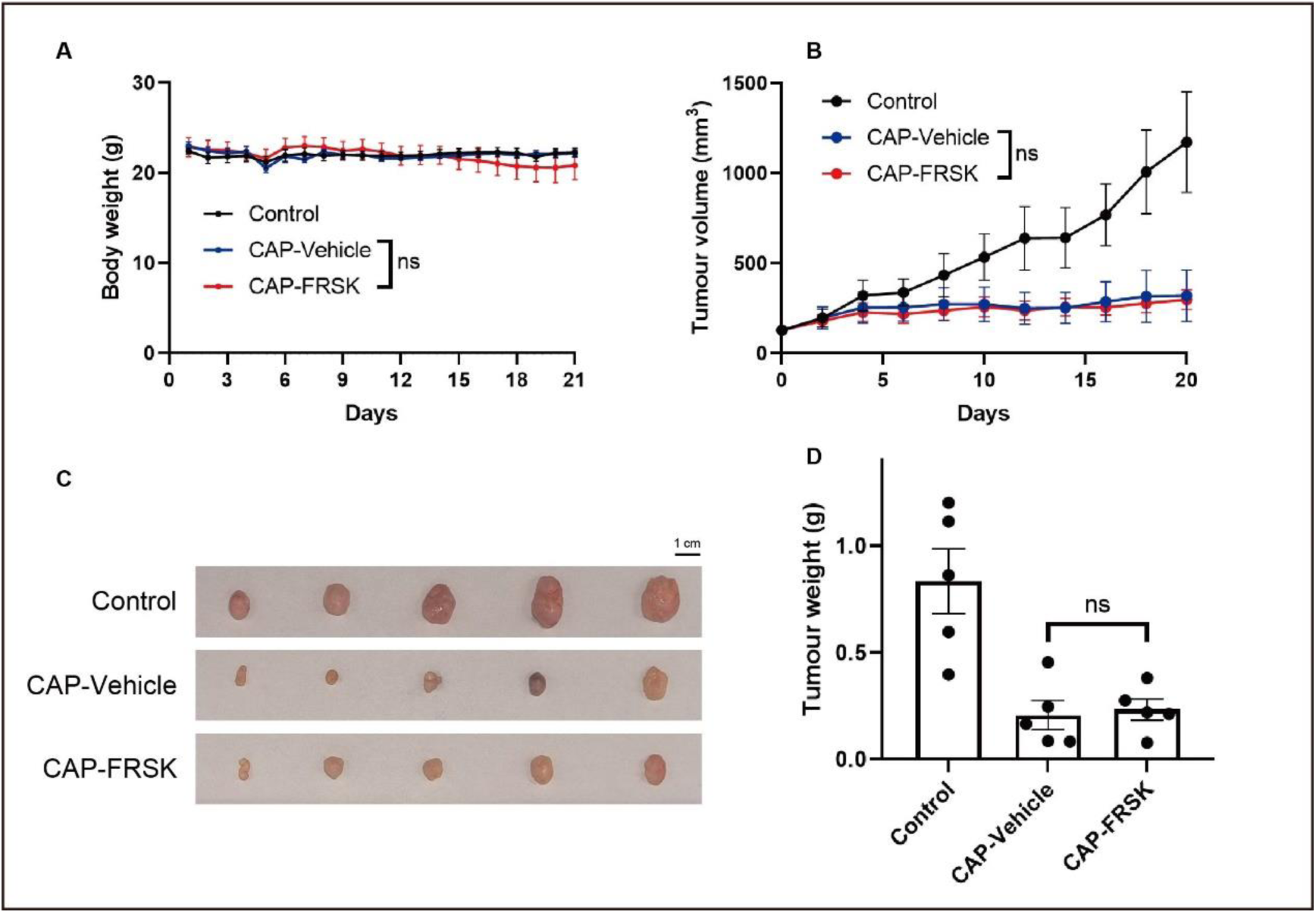
Effects of topical forskolin on anti-tumor efficacy of capecitabine. (A) Topical forskolin has no effect on body weight and (B) tumor volume change. (C) Comparison of tumor volume and (D) tumor weight.

Moreover, forskolin had no significant effect on the tumor growth inhibition induced by capecitabine **(Fig. 7B)**. By the end of the experiment, capecitabine significantly reduced tumor volume and weight, and these effects remained unchanged under forskolin treatment **(Fig. 7, C and D).**

These findings suggest that topical forskolin treatment does not impair the anti-tumor efficacy of capecitabine, supporting its potential use as a safe adjuvant drug of capecitabine.

## Discussion

This study presents a dual-filter target screening strategy that integrates literature-validated targets to refine HTS results, thereby minimizing experimental validation efforts. We applied this approach to identify targets for drugs that alleviate HFS — a painful side effect of the chemotherapy drug 5-FU — and demonstrated a marked improvement in screening efficiency. Under this dual-filter strategy, we identified ATF4 as a key target regulating 5-FU toxicity, and found that Forskolin, an ATF4 inducer, significantly reduced 5-FU toxicity and alleviated HFS.

Applying our strategy to HFS narrowed an initial list of 127 CRISPR/Cas9 screening hits down to just 2 high-confidence targets — a nearly 98% reduction in candidates requiring validation compared to conventional HTS. This dramatic streamlining of the validation process underscores the method’s efficiency in pinpointing relevant targets. Our dual-filter screening method can be broadly applied to target identification in diverse research contexts. When initial screens produce extensive candidate lists, integrating the two filtering layers (HTS-based and literature-based) helps prioritize the most promising targets. Compared to the approach introduced by Gogleva et al. (*12*), our strategy is simpler and more flexible in application, while still achieving robust target identification. However, if relevant literature is scarce, this approach may fail to flag important targets or overlook novel ones. In such cases, incorporating insights from related research fields may help compensate for gaps in the literature.

Beyond introducing a generalizable screening strategy, our study yielded clinically relevant and scientifically significant findings specific to HFS. We identified a potential new strategy for managing HFS and gained fresh insights into the mechanisms of 5-FU toxicity. Thus, this work not only streamlines target discovery but also provides a foundation for improved management of chemotherapy-induced side effects like HFS. In particular, we found that forskolin reduces 5-FU toxicity via an ATF4/eIF2α/IκB- mediated pathway. This discovery supports the view proposed by Gameiro et al. (*35*) that high levels of phosphorylated eIF2α activate NF-κB and promote inflammation. It also aligns with the well-documented anti-inflammatory effects of forskolin (*38,39*). Notably, the ATF4-driven detoxification pathway observed here differs from typical stress-induced ATF4 pathways, likely because forskolin induces ATF4 through a non-stress mechanism (*39,40*). Our findings may spur further research on the use of forskolin and inform efforts to overcome 5-FU resistance. However, because any adjuvant that reduces chemotherapy toxicity must be delivered locally to avoid compromising anti-tumor efficacy, forskolin may be suitable only for localized treatment. Future studies could focus on improving the tissue specificity of such adjuvants–potentially by leveraging additional data sources–to enable systemic administration for side-effect management without compromising anti-tumor effects, or applying specific tissue targeting delivery system.

## Materials and Methods

### 1. Genome-wide CRISPR/Cas9 screen

#### Preparation of lentiviral library

This study uses Human GeCKOv2 Library A, containing 19050 genes with 3 sgRNA per gene, 1846 microRNAs with 4 sgRNA per gene, and 1000 non-targeting control sgRNAs. Plasmid DNA was amplified by PCR. The plasmid library was packaged into lentiviruses using 293T cells. The lentivirus-containing supernatants were filtered using 0.45 μM sterile filter and save in -80℃ for subsequent experiments.

#### Library transduction

We first prepared HaCaT cells stably expressing Cas9. HaCaT cells were transduced with cas9 plasmid by lentivirus. Cells were selected until stable Cas9 expression was achieved through antibiotic treatment. Then, the plasmid library was transduced with a MOI of 0.3 into the Cas9 expressing HaCaT cells by the lentivirus-containing supernatants. The cell pools were selected with puromycin until the control cell pools were subsequently depleted.

#### Drug treatment

Following selection, cell pools are divided into three groups. Groups as follows: 1) Control group: treated with PBS; 2) IC50 group treated with 0.0024 μM 5-FU; IC99 group treated with 0.1 μM 5-FU. Cells in all groups were kept in culture for 10 days.

#### Genomic DNA extraction and sequencing

following drug treatment, the Genomic DNA was extracted separately. PCR was used to amplify the inserted sgRNAs for the next-generation sequencing.

#### Data analysis

sequence data was obtained through next-generation sequencing. In IC50 and IC99 groups, the genes targeted by at least one sgRNA account for 98% of the total genes in the library, and the genes targeted by two or more active sgRNAs comprise 85% and 87%, respectively. The sgRNA counts were analyzed by MAGeCK after the data for microRNAs been excluded. Counts were normalized based on the 1000 non-targeting control sgRNAs. RRA scores for two screen groups (Control vs IC50 and Control vs IC99) were calculated with MAGeCK. The valid genes in both screen groups were defined as the top genes that meet p-value less than 0.02 after ranking the RRA scores. The final screen outputs were defined as the genes that appears in both valid gene lists and ranked in the top 10.

### 2. Text mining screen

Data source is provided by PubMed database. The key word for searching is “5-FU AND (resistance OR sensitivity) AND (expression OR silence OR knockdown)”, and the searching scale is the title and abstract. Textual information from the scientific title and abstract was subjected to lexical analysis using an advanced tokenization algorithm, which generated normalized word tokens with corresponding vector representations through an embedding layer, BioGPT embeddings (*25*). For reference gene annotation, we compiled a comprehensive gene lexicon containing reference gene symbols and known aliases, which was subsequently transformed into a vector using equivalent embedding parameters. The original reference gene list is downloaded from NCBI, contains the gene symbol and alias of all protein-coding genes in homo sapiens. As for some replicate alias in the reference gene list, we corrected them manually. The core extraction process employed a cosine similarity-based metric (threshold ≥0.85) to quantify vector-space alignment between tokenized text elements and gene referents. GTEx database provided the gene expression data of human normal skin tissue. The expression data of the “skin_not_sun_exposed” samples were downloaded. The genes with median Transcripts Per Million (TPM) > 10 were retained. The “Intersection” program was used to integrate the outputs of “Extractor” and the retained genes from GTEx database. The outputs of “Intersection” was defined as text mining screen results.

### 3. Cell culture

HaCaT cells and 293T cells were cultured in DMEM supplemented with 10% fetal bovine serum (FBS) and 1% penicillin-streptomycin (P/S). HCT116 cells were cultured in McCoy’s 5A supplemented with 10% FBS and 1% P/S. ATF4-knockdown, ATF4- knockout, ATF4-overexpressing, and Cas9-expressing cell lines were all derived from HaCaT cells and were cultured in the same condition with Hakata. All cells were cultured in a humidified incubator at 37°C and 5% CO2.

### 4. Cell lines construction

Lentiviral particles were packaged with Lentiviral Packaging Kit (Genomeditech), incorporating the following plasmids: Human *ATF4* knockdown plasmid (shRNA- ATF4, GTTGGTCAGTCCCTCCAACAA), Human *ATF4* knockout plasmid (sgRNA- ATF4, caccg-CCACTCACCCTTGCTGTTGT), and Human *ATF4* overexpression plasmid (over-ATF4, NM_001675.4), along with their respective negative control plasmids. The lentiviruses transduction was performed on HaCaT cells for 24 hours in the presence of 8 μg/mL polybrene. Subsequently, the infected cells were selected out with 2 μg/mL puromycin for 48 hours and then cultured in DMEM containing 0.5 μg/mL puromycin for expansion. Finally, the stable HaCaT cell lines with *ATF4* knockdown, *ATF4* knockout, and *ATF4* overexpression were successfully established.

### 5. In vitro experiments

#### LDH Assay

HaCaT cells were plated at 8,000 cells per well in 96-well plates. Once adherent, cells were treated with 20 μM 5-FU together with various concentrations of forskolin. After 24 hours, the culture supernatant was collected, and lactate dehydrogenase (LDH) activity was measured according to the manufacturer’s protocol (Promega). Absorbance was recorded at 490 nm to assess cell membrane integrity and cytotoxicity.

#### CCK-8 Assay

HaCaT cells were plated at 8,000 cells per well in 96-well plates and required adherent. Cells were subsequently exposed to 20 μM 5-FU, together with gradient concentrations of forskolin. After 24 hours, CCK-8 reagent was added, and absorbance at 450 nm was measured to determine cell viability. For determination of IC50 changes, cells were exposed to a series of 5-FU concentrations for 24 hours, with or without 50 μM forskolin added. IC50 values were calculated with GraphPad Prism.

#### Cell Cycle Analysis

HaCaT cells were plated at 3×10^5^ cells per well in 6-well plates and allowed to adhere. Cells were then treated with solvents (NC group), 20 μM 5-FU (5-FU group), 50 μM forskolin (FRSK group), or a combination of 5-FU and forskolin (5-FU+FRSK group). After 24 hours, cells were collected, fixed, and stained with propidium iodide (PI). Finally, cell cycle distribution was analyzed by flow cytometry.

#### Apoptosis Assay

HaCaT cells were plated at 3×10^5^ cells per well in 6-well plates and allowed to adhere. Cells were treated with solvents, 40 μM 5-FU, 50 μM forskolin, or a combination of 5-FU and forskolin. After 12 hours, cells were collected, washed, and stained with Annexin V/PI (Beyotime) according to the manufacturer’s protocol. Finally, apoptotic and necrotic cells were quantified using flow cytometry.

#### mRNA expression analysis

HaCaT cells were plated at 3×10^5^ cells per well in 6-well plates and continually cultured 24 hours. Cells were treated with the solvent, 20 μM 5- FU, 50 μM forskolin, or a combination of 5-FU and forskolin. After 12 hours, cells were cleaned with PBS, and the lysed cells were subjected to RT-qPCR analysis.

#### Protein expression analysis

For the verification of shRNA-ATF4, sgRNA-ATF4, and over-ATF4, HaCaT cells were plated at 5×10^5^ cells per well in 6-well plates and continually cultured 24 hours. For the exploring experiments on p-eIF2α, HaCaT cells were plated at 3×10^5^ cells per well in 6-well plates and continually cultured 24 hours. HaCaT cells were treated with the solvent, 20 μM 5-FU, 50 μM forskolin, or a combination of 5-FU and forskolin for 1 hour. HaCaT cells expressing sgRNA-NC and sgRNA-ATF4 were treated with 20 μM 5-FU or a combination of 5-FU and 50 μM forskolin for 1 hour. For the exploring experiments on IκB, HaCaT cells were plated at 3×10^5^ cells per well in 6-well plates and continually cultured 24 hours. HaCaT cells expressing shRNA-NC and shRNA-ATF4 were treated with 20 μM 5-FU or a combination of 5-FU and 50 μM forskolin for 12 hours. After the drug treatment, the collected cells were subjected to Western blot analysis.

#### RNA-seq analysis

HaCaT cells expressing shRNA-NC and shRNA-ATF4 were plated at 5×10^5^ cells per well in 6-well plates and continually cultured 24 hours. After that, cells were cleaned with PBS and collected. Total RNA in the two groups were isolated and subjected to the Illumina sequencing. Differential expression analysis was performed using the DESeq2 package. Differentially expressed genes (DEGs) were defined when the absolute value of fold of change more than 2 and the p-value less than 0.05. The KEGG pathway enrichment analysis of the DEGs was performed by script in house of AZENTA.

### 6. Western blot analysis

Tissue or cell samples were lysed with RIPA buffer containing phosphatase inhibitors and 1 mM PMSF. Protein extracts were quantified using BCA protein assay kit and then heated at 100°C for 10 minutes with loading buffer. Proteins were separated by SDS-PAGE and subsequently transferred onto a PVDF membrane using the wet transfer method. The membranes were blocked at room temperature for 1 hour in 5% bovine serum albumin (BSA) prepared in Tris-buffered saline with 0.1% Tween-20 (TBST). Following blocking, the membrane was incubated with the following primary antibodies overnight at 4℃: anti-ATF4 (PTG, 10835-1-AP), anti-eIF2α (Abcam, ab169528), anti-p-eIF2α (MCE, HY-P80811), anti-MEK (PTG, 11049-1-AP), anti-p-MEK (PTG 28955-1-AP), anti-ERK1/2 (PTG, 11257-1-AP), and anti-p-ERK1/2 (PTG, 28733-1-AP). The membrane was then incubated with secondary antibodies at room temperature for 1 hour. After washed with TBST, protein bands were visualized with chemiluminescence using Super Singal West FemtoMaximum Sensitivity Substrate (Thermo Scientific). Finally, protein expression levels were evaluated by band intensity.

### 7. RT-qPCR analysis

Total RNA was extracted from HaCaT cells using a TRIzol Reagent (YaMei) according to the manufacturer’s instructions. RNA quality and concentration were determined using NanoDrop spectrophotometer. cDNA was synthesized using ReverTra Ace qPCR RT Master Mix (TOYOBO, FSQ-301) with 50 ng/μL of total RNA concentration. RT-qPCR was performed using qPCR SYBR Green Master Mix (YEASEN). Relative mRNA expression level was analyzed using 2^(-ΔΔ CT),^ with GAPDH as the internal reference gene. The primer sequences are listed in **table S6**.

### 8. Animal models

***HFS Model on SD Rats:*** female SD rats (6-8 weeks old) were randomly assigned to three groups. Rats were gavaged daily with capecitabine (4000 mg/kg per day or its vehicle (castor oil: ethanol: water, 1:1:1). After gavage, rats were restrained, and their hind paws were treated topically with either a treatment gel containing 10% forskolin or a vehicle gel. The gel was applied for 4 hours and then removed to prevent accidental ingestion. Grouped as follows: 1) Control group (n=6) gavaged with solvent and treated with vehicle gel; 2) CAP + Vehicle group (n=12) gavaged with capecitabine and treated with vehicle gel; 3) CAP + FRSK group (n=12) gavaged with capecitabine and treated with 10% forskolin gel; 4) Vehicle+FRSK group (n=3) gavaged with solvent and treated with 10% forskolin gel (only used for protein expression analysis). The progression of HFS was assessed daily based on paw skin changes by at least two researchers. The HFS model on rats could be evaluated as Grade 0 (normal), Grade 1 (slight desquamation or dry lines), and Grade 2 (peeling or erythema with slight desquamation). The Von Frey Filament test was used to test mechanical pain threshold of rats’ hind paws, recording the maximum force eliciting a withdrawal response every 7 days. The whole experiment lasted approximately 5 weeks.

***HFS Model on BALB/c Mice:*** Female BALB/c mice (6-8 weeks old) were randomly divided into three groups. Mice were gavaged daily with capecitabine (1000 mg/kg per day) or its vehicle (5% v/v gum Arabic solution, citrate buffer pH 6.0). Their hind paws were treated with either a treatment gel containing 10% forskolin or a control gel for 4 hours. Grouped as follows: 1) Control group (n=10); 2) CAP + Vehicle group (n=10); 3) CAP + FRSK group (n=10). The progression of HFS was evaluated and recorded daily as Grade 0 (normal), Grade 1 (slight dry lines), and Grade 2 (desquamation, erythema, or swelling). Paw withdrawal responses of hind paws to pinprick stimulation were assessed every 3 days. The whole experiment lasted approximately 2 weeks.

***Subcutaneous Tumor Bearing Model on Nude Mice:*** Female nude mice (6-8 weeks old) were injected subcutaneously with HCT116 human colorectal cancer cells (5×10^6^ cells) mixed with matrigel. The experiment was started as the average tumor volume reached approximately 150 mm^3^. Mice were randomly assigned to three groups and gavaged daily with capecitabine (180 mg/kg) or its vehicle (5% v/v gum Arabic solution, citrate buffer pH 6.0). After gavage, their hind paws were treated with either a treatment gel containing 10% forskolin or a control gel for 4 hours. Grouped as follows: 1) Control group (n=6); 2) CAP + Vehicle group (n=6); 3) CAP + FRSK group (n=6). Tumor length (L) and width (W) were measured every 2 days, and tumor volume (V) was calculated based on V=(L×W^2^)/2. The experiment was discontinued when tumor volume reached approximately 1500 mm^3^, and the tumors were subsequently excised, photographed, and weighed.

### 9. Histology

Rats’ hind paw skin was collected and divided into two parts. One part was subjected to western blot analysis for the detection of IκB, the other part was fixed in 4% paraformaldehyde (PFA) for at least 48 hours. The fixed tissues were embedded in paraffin and sectioned at 4 μm thickness. The sections were subjected to hematoxylin and eosin (HE) staining and immunohistochemistry (IHC) staining. The IHC staining utilized the following antibodies: CD3, CD68, Ki67. Apoptosis was evaluated using a TUNEL assay kit (Beyotime) following the manufacturer’s instructions.

### 10. Statistical analysis

All data in this study was performed with GraphPad Prism 8.0.1. The Significance of difference between two groups was analyzed by Student’s t test (unpaired, two-tailed).

For the data with more than two groups, one-way ANOVA was performed. The Error bar was presented as the mean ± standard error of mean (SEM). P<0.05 was considered as significant (ns, P>0.05; *, P<0.05; **, P<0.01; ***, P<0.001).

## Author contributions

Shiyi, Z. and Jie, L. supervised the research. Qiqi, Y. designed and performed most experiments. Bingxue, Y. aided the design of CRISPR/Cas9 screening experiment. Junkang, A. provided technical assistance in text mining. Dazhao, L. aided some animal experiments. Bingxue, Y. and Shuyue, X. performed some in vitro experiments. Qiqi, Y. analyzed and visualized the data. Qiqi, Y. drafted the manuscript. Shiyi, Z., Jie, L., and Qing, Y. reviewed and edited the manuscript. All authors discussed and approved the manuscript. The authors declare no competing financial interest.

## Acknowledgments

We thank Ju Zeng and Heyuan Xiao for insightful discussions and manuscript editing. We are thankful to the Shanghai Cancer Institute for providing the technical platform. Finally, we acknowledge the contributions of all animals used in this study.

## Notes

### Competing Interest Statement

The authors have declared no competing interest.

### Summary of Updates

methods and materials updated; figures updated; author contribution updated.

