## Supplementary materials for "Integrating Literature Evidence to Enhance CRISPR-based Target Screening for Hand-Foot Syndrome" (from another version of this preprint: version 1, 2025-03-13)

Qiqi Yang *et al.*

**This PDF file includes:**

Figs. S1 to S5
Legends for Tables S1 to S5
Table S6

**Other Supplementary Materials for this manuscript include the following:**

Tables S1 to S5

**Supplemental Figures**

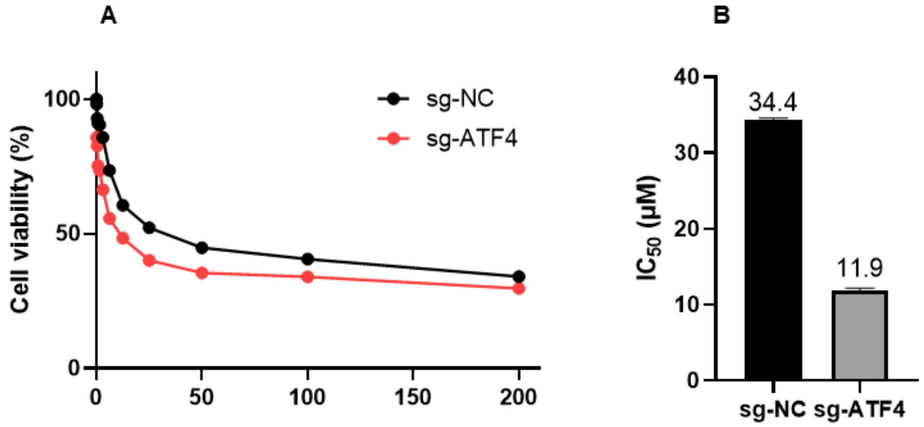


**Fig. S1. Related to Fig. 3. Effects of ATF4 knockout on 5-FU toxicity of HaCaT.**

(A) Cell viability changes after ATF4 knockout.

(B) $IC_{50}$ changes after ATF4 knockout.

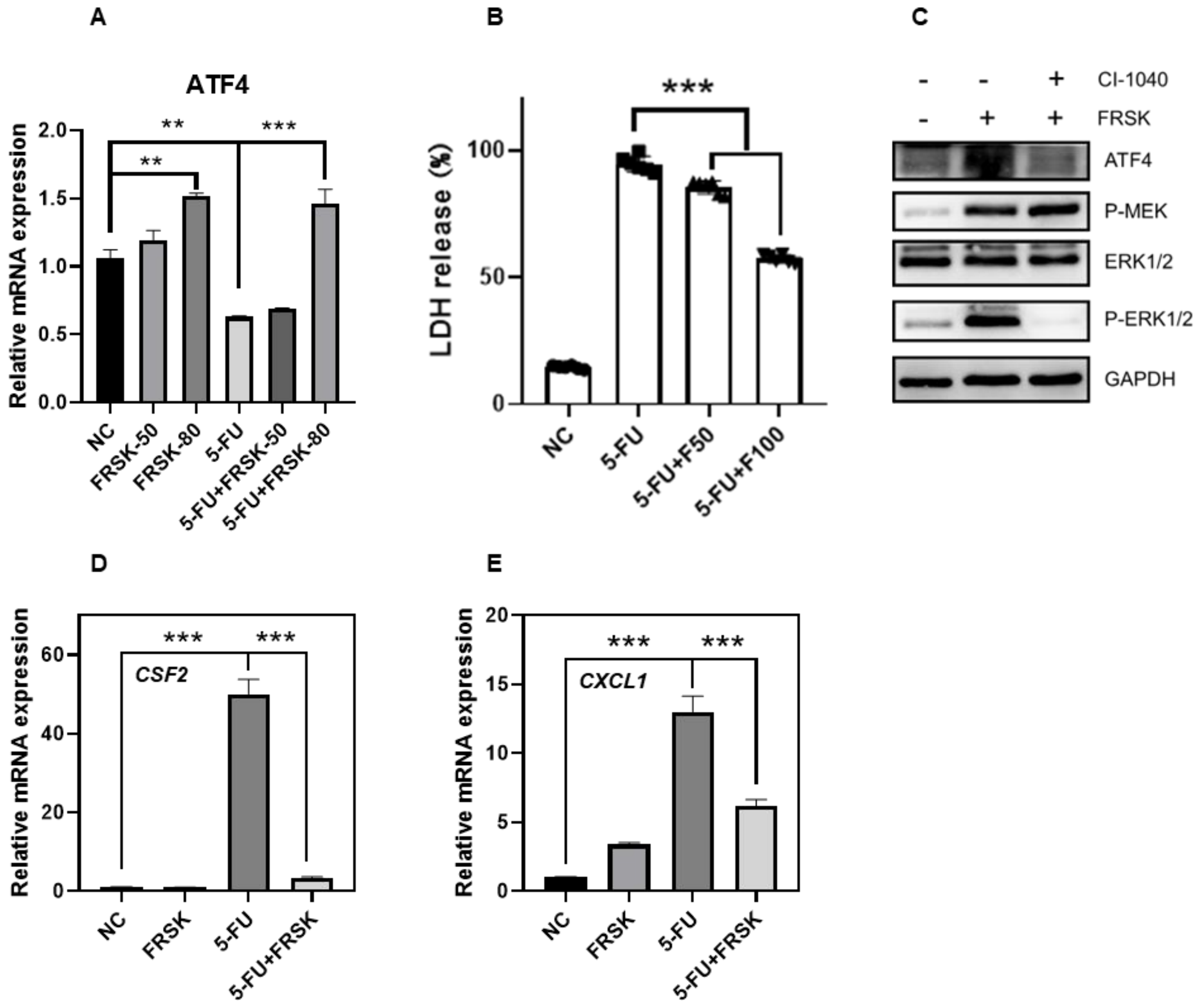


**Fig. S2. Related to Fig. 4. In vitro evaluation for the ability on detoxifying 5-FU of forskolin.**

(A) Effects of forskolin on relative mRNA expression of *ATF4*.
(B) Effects of forskolin on 5-FU-induced LDH release.
(C) Effects of MEK inhibitor on ATF4 protein expression after forskolin treatment.
(D)–(E) Effects of forskolin on 5-FU-upregulated *CSF2* and *CXCL1*.

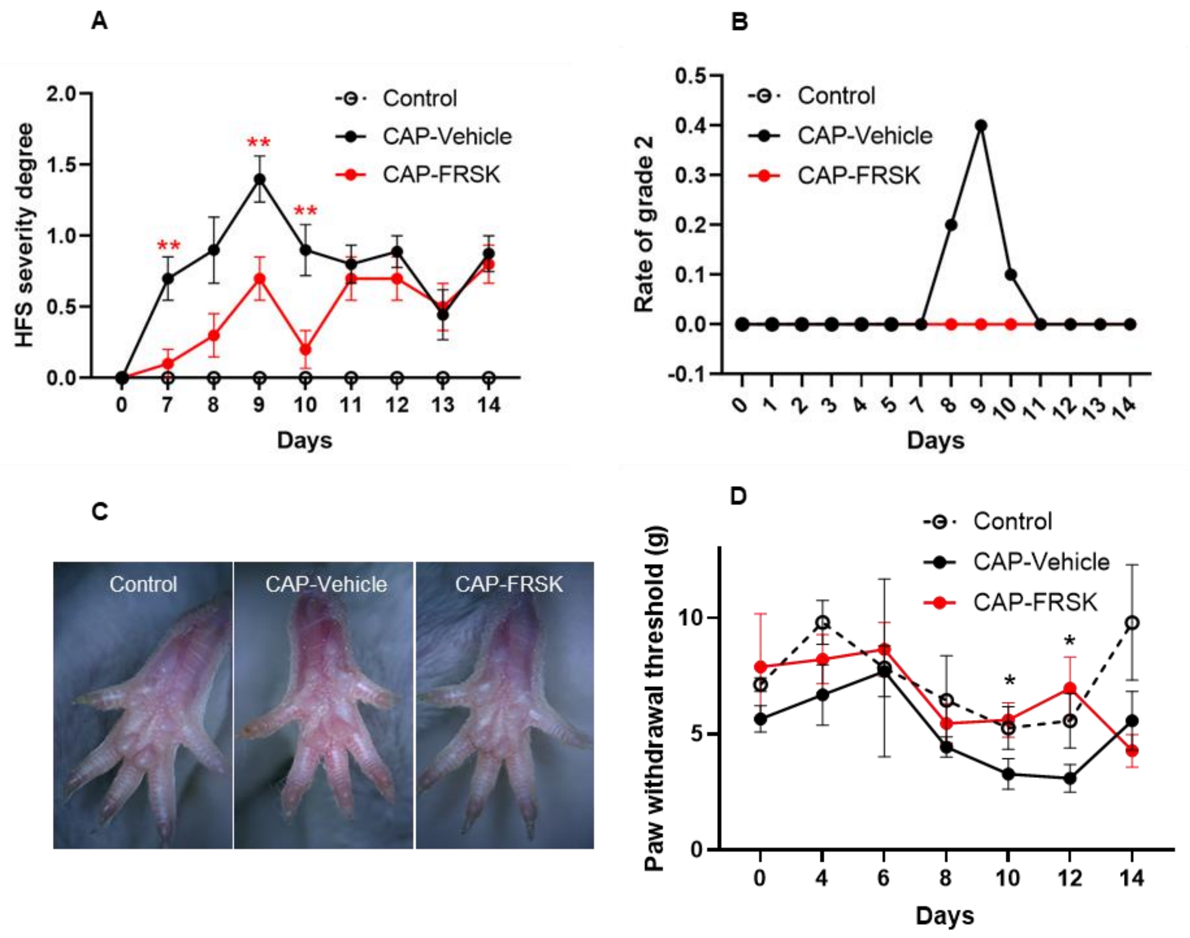


**Fig. S3. Related to Fig. 5. Efficacy of forskolin in alleviating HFS evaluated on Babl/c mice.**

(A) Effects of forskolin on capecitabine-mediated HFS severity degree evaluated on Babl/c mice.

(B) Effects of forskolin on rate of grade 2 HFS induced by capecitabine.

(C) Representative images of HFS symptoms on the hind paw of Babl/c mice.

(D) Effects of forskolin on paw withdrawal threshold.

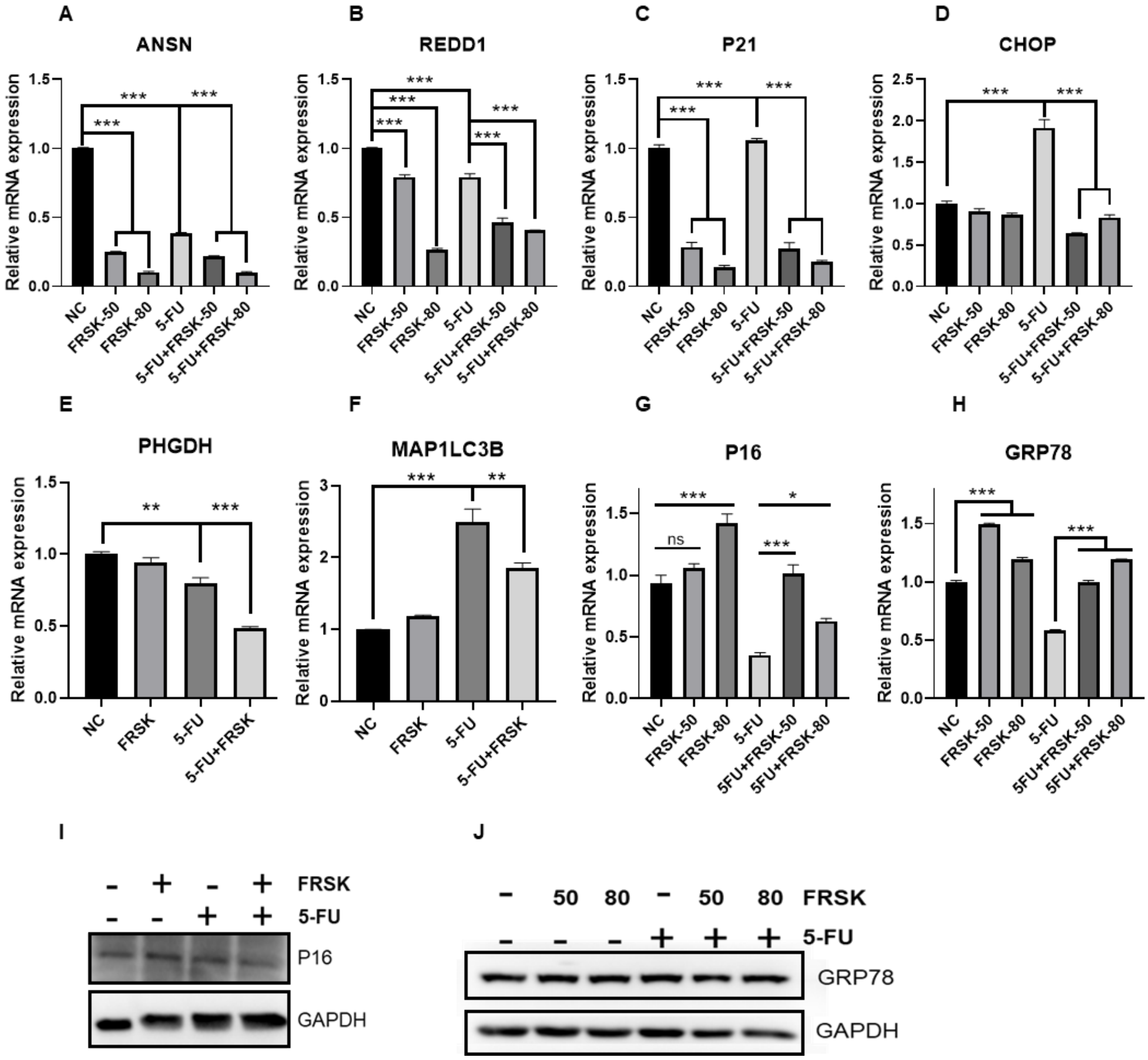


**Fig. S4. Related to Fig. 6. Effects of forskolin on the downstream stress-response effectors of ATF4.**

(A)-(H) Effects of forskolin on the relative mRNA expression of *ANSN*, *REDD1*, *P21*, *CHOP*, *PHGDH*, *MAP1LC3B*, *P16*, *GRP78*.

(I), (J) Effects of forskolin on the protein expression of P16 and GRP78.

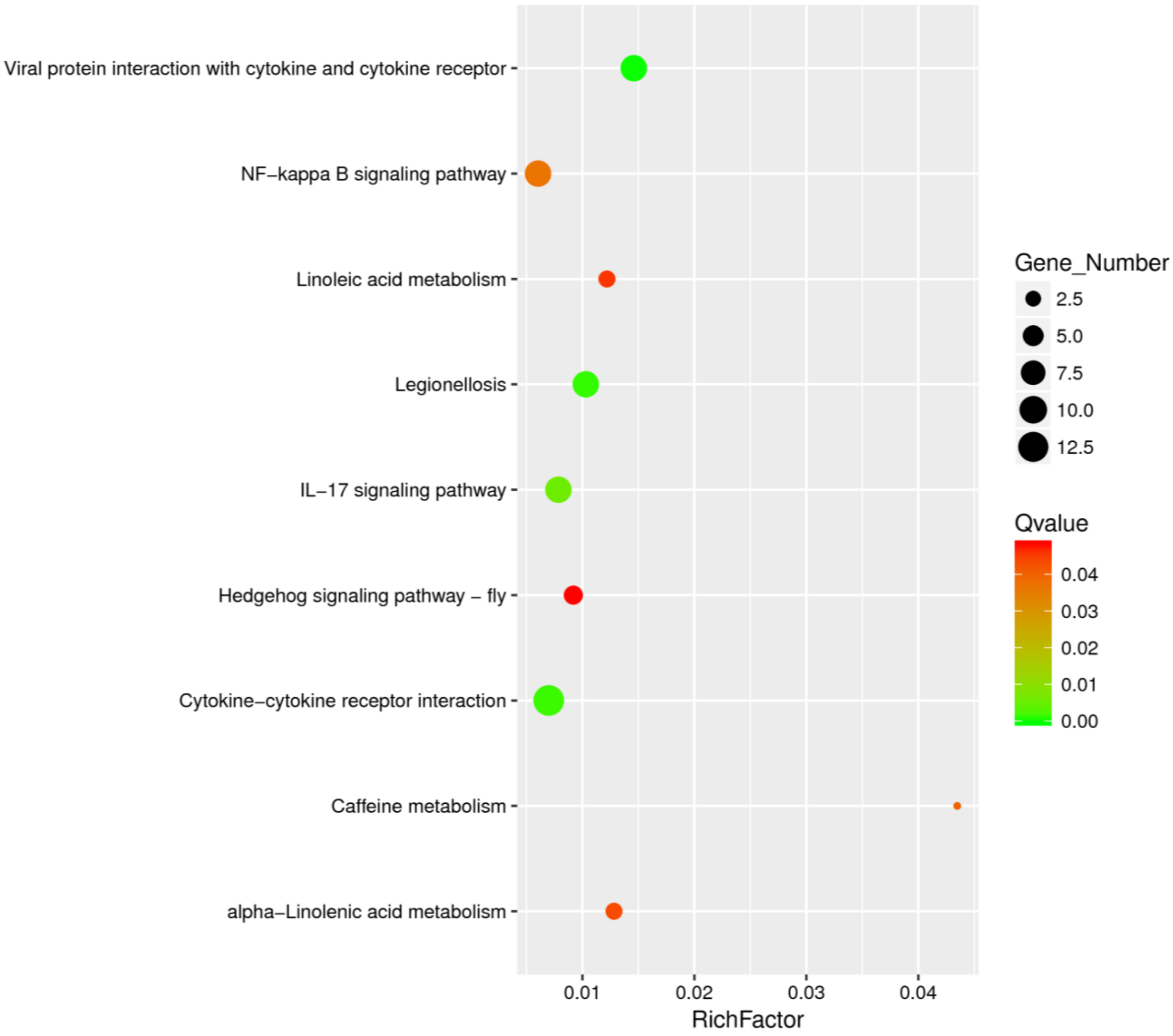


**Fig. S5. Related to Fig. 6. KEGG enrichment analysis of DEGs produced by *ATF4* knockdown.**

**Supplemental Tables**

**Table S1. Related to Fig. 1. Normalized sgRNA counts from CRISPR/Cas9 screening. (separate file)**

**Table S2. Related to Fig. 1. MAGeCK analysis results of CRISPR/Cas9 screen data. (separate file)**

**Table S3. Related to Fig. 1. Definition of targets library from CRISPR-Cas9 screening. (separate file)**

**Table S4. Related to Fig. 2. Reference list of gene symbol and alias from NCBI database. (separate file)**

**Table S5. Related to Fig. 2. Targets library from text mining. (separate file)**

**Table S6. Sequences of primers used for RT-qPCR.**

| Genes | Forward (5'-3') | Reverse (5'-3') |
|---|---|---|
| *GAPDH* | TGCCCTCAACGACCACTTTG | TTCCTCTTGTGCTCTTGCTGGG |
| *TNFA* | CTGGTCCCCAAAAGAAATGGAG | CTGGGCCACTGACTGATTTGTG |
| *IL1B* | CGCCAGTGAAATGATGGCTTATTAC | GTCGGAGATTCGTAGCTGGATG |
| *IL6* | AACAACCTGAACCTTCCAAAGATG | GCTTGTTCCTCACTACTCTCAAAT |
| *CSF2* | CACTGCTGCTGAGATGAATGAAA | GTCTGTAGGCAGGTCGGCTC |
| *CXCL1* | GCTTGCCTCAATCCTGCATC | AGTTGGATTTGTCACTGTTCAGC |
| *HSPA5* | TGACCAGAATCGCCTGACAC | TGTCAGCATCTTGGTGGCTT |
| *P16* | CACCGAATAGTTACGGTCGG | GCACGGGTCGGGTGAGAGTG |
| *NRF2* | *ACATCGAGAGCCCAGTCTTC* | *AGCTCCTCCCAAACTTGCTC* |
| *MAP1LC3B* | *TCGAACAAAGAGTAGAAGATGTCCG* | *AGCATTGAGCTGTAAGCGCC* |
| *PSAT1* | *AAAAACAATGGAGGTGCCGC* | *GGCTCCACTGGACAAACGTA* |
| *PHGDH* | *CCAATGGGAACAGCCTCAGT* | *ACTGGGCAAAGGTGTTGTCA* |
| *CHOP* | *TTCTCTGGCTTGGCTGACTG* | *CTGCGTATGTGGGATTGAGG* |
| *ATF4* | *CCCCTTCACCTTCTTACAACC* | *GGGCTCATACAGATGCCACTA* |
| *ANSN* | *GCAGCTGAAAGAAGCCCAAGT* | *TGTCTTCCATGCCAATTGCA* |
| *REDD1* | *TGGTGCCCACCTTCCAGCTG* | *GTCAGGGACTGGCTGAAGCC* |
| *P21* | *CCTGTCACTGTCTTGTACCCTTG* | *TGGTAGAAATCTGTCATGCTGGTC* |
